# Testing different forms of regulation of yolk thyroid hormone transfer in pied flycatchers

**DOI:** 10.1101/2020.04.23.056994

**Authors:** Tom Sarraude, Bin-Yan Hsu, Ton G.G. Groothuis, Suvi Ruuskanen

**Author notes:** Corresponding author: Tom Sarraude.

## Abstract

Hormones transferred from mothers to their offspring are considered a maternal tool to prepare progeny for expected environmental conditions, increasing maternal fitness. To flexibly influence offspring, mothers should be able to transmit the hormonal signals independent of their own hormonal status. However, the ability to regulate hormone transfer to the next generation is under debate. We studied the transfer of thyroid hormones (THs) to eggs in a bird model. We elevated thyroxine (T_4,_ the prohormone for the biological active triiodothyronine, T_3_) during egg-laying using T_4_ implants on females of a wild population of pied flycatchers (*Ficedula hypoleuca*), and measured plasma and yolk T_4_ and T_3_ as a response. To our knowledge, studies that manipulated a prohormone and measured the change in its active metabolites have rarely been conducted. We found an increase in plasma and yolk T_4_ and no change in T_3_ concentrations leading to a similar decrease in yolk T_3_/T_4_ ratio in response to the T_4_ treatment in plasma and yolk. This suggests that mothers are able to regulate the conversion of T_4_ in T_3_ to avoid potential costs of elevated exposure to the active hormone to herself and to her progeny. Finally, contrary to our predictions, we found no evidence of regulatory mechanisms at the follicle level, which is essential for independent regulation of yolk hormone transfer.

**Summary statement:** Thyroid hormones have been overlooked in the context of hormone-mediated maternal effects. We found that mothers may regulate yolk thyroid hormone transfer by regulating the conversion of the active form of the hormone.

## Introduction

Maternal effects are the non-genetic influences of a mother on her progeny and are thought to be adaptive (Moore et al., 2019; Mousseau and Fox, 1998; Yin et al., 2019). Maternal hormones transferred to the next generation are a potential prenatal pathway for mothers to shape their offspring phenotype (Groothuis et al., 2005; Groothuis et al., 2019; Ruuskanen and Hsu, 2018, 20). Mothers transfer thyroid hormones (THs) that have so far received little attention compared to glucocorticoids and androgens (Ruuskanen and Hsu, 2018). THs are produced by the thyroid gland and are present in two main forms: thyroxine (T_4_) and triiodothyronine (T_3_). T_4_, a precursor of T_3,_ is converted to T_3_ in tissues. T_3_ has a much greater affinity to TH receptors than T_4_. Thyroid hormones have pleiotropic effects that serve several biologically important functions across vertebrates, including growth, reproduction and metabolism (Ruuskanen and Hsu, 2018).

For maternal hormone transfer to be adaptive, mothers should be able to regulate their deposition according to the expected environment during the offspring development. Regulatory mechanisms of maternal hormone transfer are also essential to minimise physiological trade-offs between optimal hormone exposure for the mother versus that for the offspring (Groothuis and Schwabl, 2008). The evidence for a regulatory mechanism for several hormones, including corticosterone and THs is mixed (Groothuis and Schwabl, 2008), but such regulation could take place at the circulating level in the mothers and/or at the follicle level. Regulation at the follicle level may happen by controlling the transfer or conversion of THs or by producing THs independently from the thyroid gland. These mechanisms have been suggested to exist in human ovaries (Monteleone et al., 2017; Rae et al., 2007). If such regulatory mechanisms exist in other taxa, mothers may be able to regulate the deposition of THs in their eggs independently from their own circulating TH levels. This would free mothers from the possible constraint to optimise their own circulating levels of THs and the levels in their eggs independently of each other. A few studies in birds have shown some preliminary evidence that mothers may indeed be able to regulate yolk TH transfer. In Japanese quails, a low-dose oral administration of T_4_ resulted in an increase in yolk T_3_ but not in circulating T_3_, whereas T_4_ increased in both tissues (Wilson and McNabb, 1997). Administration of T_3_ in turn increased plasma T_3_ but not yolk T_3_ (Wilson and McNabb, 1997). Furthermore, artificial blocking of TH production in hens led to a decrease in yolk T_3_ but not in plasma T_3_, while T_4_ decreased in both tissues (Van Herck et al., 2013). These studies on captive birds induced long exposure to elevated hormones and, importantly, do not provide clear evidence for any of the regulatory mechanisms introduced above. Therefore, there is a need for complementary studies under shorter time scales relevant for wild passerine species.

In this experiment we tested whether mothers are able to regulate their transfer of yolk THs, at the circulating and/or at the follicle level. We experimentally manipulated TH levels with T_4_ implants using a within-subject design in female pied flycatchers (*Ficedula hypoleuca*) during egg-laying and collected plasma samples and pre-and post-implantation eggs for the analysis of T_3_ and T_4_. Implanting the prohormone T_4_ would enable us to test the potential differential conversion of this hormone to the biological active T_3_ in the mother as a regulatory mechanism to protect the egg from increased exposure to these hormones. It would also test whether mothers can regulate the transfer of hormones to the egg. First, if the implant successfully increased circulating T_4_, we would expect an increase in circulating T_3_ as well due to higher availability of the substrate (i.e. the prohormone T_4_) and conversion to T_3_. Alternatively, females may buffer the increase in plasma T_4_ by downregulating the conversion of T_4_ to T_3_, thus yielding no increase in plasma T_3_. Second, we predict that if mothers were able to regulate yolk TH transfer independently from their circulating levels, only one of these two compartments (i.e. plasma or yolk) would be affected by exogenous T_4_, or one would be more affected than the other (Groothuis and Schwabl, 2008). In this case, the T_3_/T_4_ ratio may be different between the two tissues. Conversely, if mothers were unable to regulate yolk TH transfer, one would expect both plasma and yolk THs to vary in the same direction and with a similar magnitude in response to exogenous T_4_ (Groothuis and Schwabl, 2008). Thus, the T_3_/T_4_ ratio would not differ between the tissues.

## Material and methods

The experiment was conducted in 2016 and 2017 in Turku, Finland (60°26’N, 22°10’E). The study species, the pied flycatcher, generally lays a single clutch of 6 to 7 eggs. We inserted in egg-laying females either a T_4_ implant (10 μg, hereafter T4) or a control implant (hereafter CO) (Innovative Research America, Sarasota, FL, USA). The amount of T_4_ was aimed to mimic natural T_4_ production and was designed to release the hormone steadily over 21 days (see below for more details on the dose and implantation).

### Preparation of T_4_ implants and implantation

Two types of sterile T_4_ implants (ca. 3 mm of diameter) were used for this experiment: ready-made T_4_ pellets (10 μg, hereafter PT4) and respective controls (PCO) which were identical, but without T_4_. Both implants were produced by Innovative Research America (Sarasota, FL, USA). The amount of T_4_ in the implants was based on the production rate of T_4_ measured in chickens, quail and pigeons (1–3 μg T_4_/100 g of body weight per day (McNabb and Darras, 2015) and adjusted to the average body mass of pied flycatchers. T_4_ is embedded in a matrix that is designed to steadily release the hormone for 21 days.

Before implantation between the scapula, the skin was disinfected with a cotton pad dipped in 70% ethanol. An incision was made with a 18G needle (BD Microlance ™) and the implant was inserted and pushed away from the incision to avoid losing the implant. The wound was sealed with veterinary tissue adhesive 3M Vetbond™, commonly used in experiments with pit tags and shown to have no effects on birds.

### Experimental design - captive females

First, to validate that implants increased circulating THs in a short time window after implantation, we conducted an experiment with female pied flycatchers in captivity. Since the yolk formation takes approximately 3.5 to 4 days in passerines (Williams, 2012), implants inserted during egg-laying need to increase hormone levels within days to be able to quantify their effect on newly formed eggs. Captive birds (on natural photoperiod and ad libitum food) were used for the validation experiment as repeated disturbance during egg-laying in the wild population could have caused nest desertion. On the 4^th^ day after capture, each female received either a subcutaneous control or T_4_ implant (n = 4 per group). Blood samples were taken before the implant was inserted, 24 hours and 72 hours after the implant (between 09.30-11.30 a.m.). Circulating TH levels in response to the implants are presented in Table 1.

**Table 1:** Circulating T_4_ and T_3_ in captive female pied flycatchers in response to T_4_ (T4) or control (CO) implants. Females were sampled prior to insertion of the implant, 24h and 72h later.

| Implant | CO |  |  | T4 |  |  |
| --- | --- | --- | --- | --- | --- | --- |
| Hour | 0 | 24 | 72 | 0 | 24 | 72 |
| N | 3 | 4 | 4 | 3 | 3 | 3 |
| Mean (SD) |  |  |  |  |  |  |
| $T_4$ (in pg/μl) | 4.9<br>(1.7) | 3.5<br>(1.3) | 4.9<br>(1.9) | 5.3<br>(2.6) | 28.9<br>(18.0) | 8.4<br>(3.3) |
| Mean (SD) |  |  |  |  |  |  |
| $T_3$ (in pg/μl) | 0.7<br>(0.5) | 0.5<br>(0.4) | 0.9<br>(0.5) | 0.7<br>(0.1) | 0.7<br>(0.2) | 1.1<br>(0.2) |

### Experimental design - wild females

The first egg of a clutch was collected and replaced by a dummy egg as a within-clutch control (hereafter “pre-implant”). On the morning of the second egg (7–9 a.m.), females were captured and received a T_4_ or a control implant as above. The last egg of a clutch (mean egg rank (SD) = 6 (0.25), hereafter “post-implant”) was collected, as it is mostly formed under the influence of the hormone implant (see above). Early in the incubation, on average 1.3 days (SD = 1.0) after the last egg laid, females were blood sampled for the analysis of circulating T_4_ and T_3_.

### Hormone analysis

Blood samples (ca. 40 μl) were taken from the brachial vein. Plasma was collected via centrifuging and frozen at −20°C until analysis. Eggs were thawed, yolks separated, homogenised in MilliQ water (1:1) and a small sample (ca. 50 mg) was used for TH analysis. Yolk and plasma THs were analysed using nano-LC-MS/MS, following Ruuskanen et al. (2018; 2019). TH concentrations, corrected for extraction efficiency, are expressed as pg/mg yolk or pg/ml plasma.

### Statistical analysis

Data were analysed with the software R version 3.6.2 (R Core Team, 2020). Linear mixed models (LMMs) were fitted using the R package *lme4* (Bates et al., 2015). P-values were obtained by model comparison using Kenward-Roger approximation from the package *pbkrtest* (Halekoh and Højsgaard, 2014). Estimated marginal means and standard errors (EMMs ± SE) were derived from models using the package *emmeans* (Lenth, 2019). Effect size estimates (Cohen’s *d)* obtained from marginal means were computed with the package *emmeans*. Effect size estimates obtained from the raw data were calculated with the package *effsize* (Torchiano, 2020). Model residuals were checked for normality and homogeneity by visual inspection.

Yolk THs were log-transformed to achieve normal distribution of the residuals. Yolk TH concentrations and T_3_/T_4_ ratio were analysed by fitting linear mixed models that included the treatment (i.e., T_4_ or CO implant) as the predictor, hormone levels in the pre-implant egg and year as covariates, and the hormone assay as a random intercept.

Plasma TH levels of the incubating females were analysed using linear regressions with the type of implant as a fixed factor and body mass, ambient temperature and time of the day as covariates, as these covariates are known to influence circulating levels (McNabb and Darras, 2015). Covariates were centred and scaled. Year was not included in the model as it covaried with ambient temperature (VIF > 2), and the latter is known to affect circulating THs (McNabb and Darras, 2015).

Effect sizes (Cohen’s *d*) of the treatment on yolk and wild female plasma THs were estimated from marginal means. Effect size estimates of the treatment on the T_3_/T_4_ ratio in the yolk and in captive female plasma were computed from the raw data. To avoid nest abandonment, we did not blood sample wild females during egg laying. Therefore, we used the data from captive birds in the following way: Plasma samples from captive females averaged over day 1 and 3 after the implantation (reflecting the yolking phase of the last egg in wild birds) were compared with the post-implant last eggs collected from wild females.

### Ethical note

The experiments were conducted under licenses from the Animal Experiment Board of the Administrative Agency of South Finland (ESAVI1018/04.10.07/2016) and South-Western Finland Centre for Economic Development, Transport and Environment (VARELY/412/2016).

## Results

Yolk THs of pre-implant eggs (first eggs of a clutch) did not differ between females with CO or T_4_ implants (mean yolk T_4_ (SE), CO = 8.29 (0.61) vs T4 = 7.63 (0.43); mean yolk T_3_ (SE), CO = 2.85 (0.20) vs T4 = 2.94 (0.38); all *t* ≤ 0.89 and all p ≥ 0.39). After receiving a T_4_ implant, females produced eggs with ca. two times higher yolk T_4_ concentration than CO-implanted females (Estimated marginal means, EMMs ± SE: post-implant treated egg = 17.14 ± 2.07 pg/mg yolk vs post-implant control egg = 8.54 ± 1.08 pg/mg yolk, Table 2, Figs. 1A, 2A). However, post-implant yolk T_3_ did not differ between the groups (mean (SE), CO = 2.34 (0.27) vs T4 = 2.72 (0.29); Table 2, Figs. 1B, 2A).

**Table 2:** (a) Full linear mixed models of yolk THs in response to T_4_ implants, in pied flycatchers (sample sizes: T4 = 13 eggs; CO = 11 eggs). Hormone assay was included as a random intercept. (b) Full linear models of plasma THs in response to T_4_ implants (sample sizes; T4 = 11 females; CO = 10 females). Covariates were centred and scaled. Ndf =1. Significant p values are shown in bold.

| (a) | Estimate (SE) | $F_{\text{ddf}}$ | P value |
| --- | --- | --- | --- |
| **Yolk T<sub>4</sub>** |  |  |  |
| Implant (T4) | 0.75 (0.19) | 14.96<sub>17.6</sub> | **0.001** |
| Pre-implant egg | 0.07 (0.06) | 1.09<sub>20.0</sub> | 0.31 |
| Year (2017) | 0.09 (0.20) | 0.18<sub>18.2</sub> | 0.67 |
| **Yolk T<sub>3</sub>** |  |  |  |
| Implant (T4) | 0.12 (0.11) | 1.09<sub>17.1</sub> | 0.31 |
| Pre-implant egg | 0.20 (0.05) | 13.33<sub>18.1</sub> | **0.002** |
| Year (2017) | -0.01 (0.12) | 0.01<sub>17.8</sub> | 0.92 |
| **T<sub>3</sub>/T<sub>4</sub> ratio** |  |  |  |
| Implant (T4) | -0.12 (0.03) | 12.28<sub>17.6</sub> | **0.003** |
| Pre-implant egg | -0.01 (0.11) | 0.01<sub>19.3</sub> | 0.91 |
| Year (2017) | -0.01 (0.03) | 0.10<sub>18.2</sub> | 0.76 |
| (b) | Estimate (SE) | $t$ | P value |
| **Plasma T<sub>4</sub>** |  |  |  |
| Implant (T4) | 1.16 (0.75) | 1.54 | 0.14 |
| Body mass | -0.90 (0.38) | -2.33 | **0.03** |
| Temperature | 0.11 (0.39) | 0.29 | 0.78 |
| Time | -0.46 (0.39) | -1.80 | 0.26 |
| **Plasma T<sub>3</sub>** |  |  |  |
| Implant (T4) | 0.14 (0.15) | 0.95 | 0.36 |
| Body mass | -0.02 (0.08) | -0.26 | 0.80 |
| Temperature | -0.05 (0.08) | -0.26 | 0.58 |
| Time | -0.09 (0.08) | -1.16 | 0.26 |

**Figure 1:**
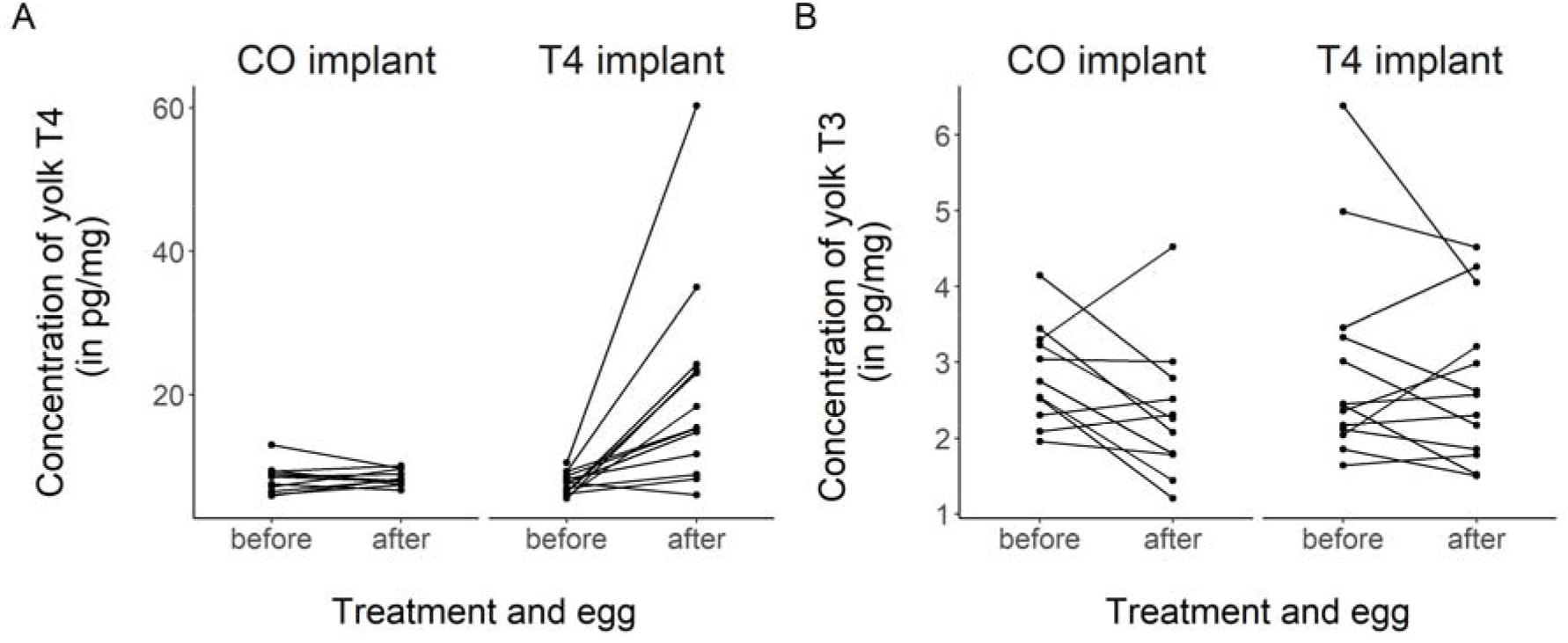
Concentrations of T_4_ (A) and T_3_ (B) in eggs of female pied flycatchers implanted with a control implant (CO implant), or 10 μg T_4_ implant (T4 implant). “Before” and “after” respectively refer to eggs collected before or after the females had received an implant.

**Figure 2:**
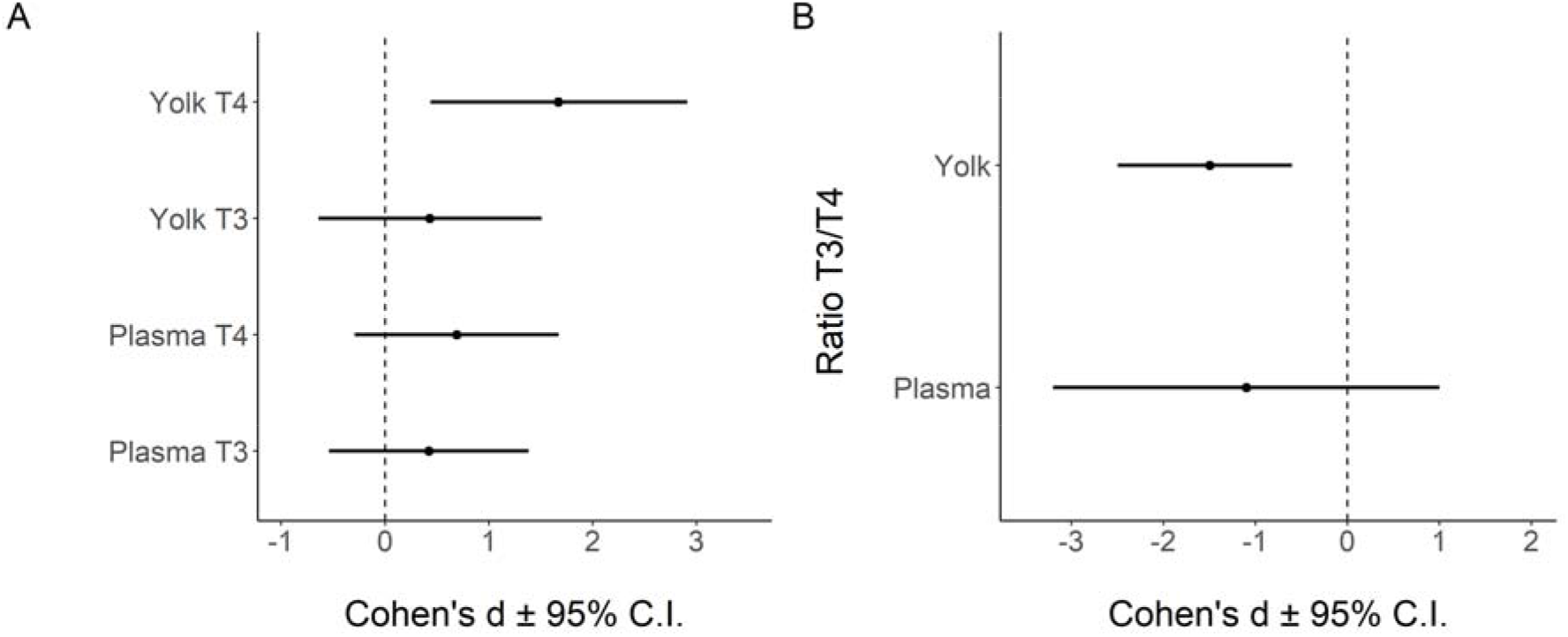
(A) Cohen’s *d* and 95% CIs for yolk (post-implant egg) and plasma (wild females) T_3_ and T_4_ calculated from the marginal means of the respective models. (B) Cohen’s *d* and 95% CIs for the T_3_/T_4_ ratio in the yolk (post-implant egg) and plasma (captive females) calculated from the raw data.

Regarding circulating THs, captive females implanted with a T_4_ implant had higher circulating T_4_ than captive control females during the first 3 days after the implant (Table 1). There was a similar, but non-significant, trend in wild female plasma T_4_ early in the incubation (when egg laying was finished, so about 10 days after implantation) (Table 2). Plasma T_3_ was not affected by the implants (Tables 1, 2).

The treatment decreased the T_3_/T_4_ ratio in the post-implant egg (Table 2) and the trend was similar for plasma levels (Fig. 2B), and a visual inspection of the effect sizes for T_3_/T_4_ ratios showed no difference between the two tissues (Fig. 2B).

## Discussion

To our knowledge, this study is the first one to manipulate circulating thyroid hormones (THs) of a wild bird species during egg-laying, to study potential regulation of maternal TH transfer at the level of mothers’ circulation and at the follicle level. Contrary to previous studies on other maternal hormones (e.g. steroid hormones), we not only looked at the response in the implanted hormone T_4_, but also at its active metabolite T_3_. We found an increase in plasma and yolk T_4_ in response to exogenous T_4_ but not in T_3_, while the T_3_/T_4_ ratio did not appear to differ between the tissues. This result would indicate an absence of regulation of transfer of these two hormones from mothers to eggs (supporting the epiphenomenon hypothesis in Groothuis and Schwabl (2008). We predicted that elevated plasma T_4_ would increase plasma or yolk T_3_, the more potent hormone, because of the increased amount of its precursor, T_4_. However, we observed no changes in circulating or yolk T_3_. This result, together with the rapid decrease in plasma T_4_ after implantation observed in captive females, suggest a change in the peripheral TH metabolism to quickly remove excess T_4_. In rats, hyperthyroidism increases the conversion of T_4_ and T_3_ into inactive metabolites (Bianco et al., 2002). Likewise, increased circulating T_4_ rapidly decreases the conversion of T_4_ into T_3_ (Bianco et al., 2002). Both mechanisms prevent the production of T_3_, which may explain why we observed no increase in plasma T_3_. These mechanisms may allow individuals to cope with elevated THs and could be important tools for mothers to protect themselves and their progeny from the potentially detrimental consequences of elevated T_3_. To ascertain this hypothesis, one should analyse the expression and activity of different enzymes involved in TH metabolism in response to exogenous THs, both in mothers and in embryos.

In addition to regulating their own plasma levels of THs, we hypothesised that mothers may be able to regulate the exposure of the developing follicles to THs. We found no evidence for such regulatory mechanism, as the T_3_/T_4_ ratio appeared not to differ between female plasma and yolk. This result is contrary to that of Wilson and McNabb (Wilson and McNabb, 1997), where yolk T_3_, but not circulating T_3_ was increased in response to long-term T_4_ administration. This contradiction may be caused by different time scales between the two studies. In our study, the peak in T_4_ rapidly decreased after implantation. Conversely, Wilson and McNabb administrated exogenous T_4_ for longer periods of time, which might have forced females to deposit T_3_ in their eggs to maintain normal plasma T_3_.

We found that females regulated the concentrations of plasma T_4_ and T_3_ but not TH transfer to the yolk. Whether the first regulatory mechanism has been selected to benefit the mother or the offspring is yet unclear. This could be tested by elevating plasma and yolk T_3_ and measuring whether potential detrimental effects are larger in the mother or the offspring. Besides, further studies could also aim at investigating the changes in thyroid hormone metabolism (enzyme production and activity) in response to increased hormones.

## Acknowledgements

We thank Sophie Michon and Florine Ceccantini for her help on the field. Mass spectrometry were performed at the Turku Proteomics Facility, University of Turku, supported by Biocenter Finland.

## Data accessibility

Data have been deposited on the online repository Zenodo. DOI: 10.5281/zenodo.3747401

## Authors’ contribution

TS and SR designed the study and collected the data. TS analysed the data and wrote the first draft of the manuscript. SR analysed TH concentrations. All authors edited the manuscript and approved its final version.

